# Comparative analysis of protein expression between oesophageal adenocarcinoma and normal adjacent tissue

**DOI:** 10.1101/2024.09.20.614054

**Authors:** Ben Nicholas, Alistair Bailey, Katy J McCann, Robert C. Walker, Peter Johnson, Tim Elliott, Tim J. Underwood, Paul Skipp

## Abstract

Oesophageal adenocarcinoma (OAC) is the 7th most common cancer in the United Kingdom and remains a significant health challenge. This study presents a proteomic analysis of seven OAC donors complementing our previous neoantigen identification study of their human leukocyte antigen (HLA) immunopeptidomes. Using label-free mass spectrometry proteomics, we compared OAC tumour tissue to matched normal adjacent tissue (NAT) to identify differentially expressed proteins.

We identified differential expression of a number of proteins previously linked to OAC and other cancers. We observed enrichment of processes and pathways relating RNA processing and the immune system. Our findings also offer insight into the role of the protein stability in the generation of the neoantigen we previously identified. These results provide independent corroboration of existing oesophageal adenocarcinoma biomarker studies that may inform future diagnostic and therapeutic research.

## Introduction

Oesophageal adenocarcinoma (OAC) accounts for about 2% of all cancer diagnosis in the UK, with an increase in 10-year survival from 4% to 12% in the last 50 years [1]. Treatment options centre on resection of the oesophagus in early-stage OAC, and chemo-or radiotherapy combined with surgery for later stage OAC [2]. Previously we presented proof-of-concept findings using mass spectrometry proteomics to identify human leukocyte antigen (HLA) presented neoantigens from a cohort of OAC donors as targets for cancer vaccines [3]. Here we present, from 7 of our OAC donors, a comparative proteomic analysis of OAC tumour tissue to matched normal adjacent tissue (NAT) (Table S1). Using label free quantification (LFQ) of bottom-up mass spectrometery proteomics we sought to identify differential expression of proteins (DEP) between OAC and NAT that may inform diagnosis and treatment options.

## Results

We quantified 3552 proteins across 7 patients using label free quantification (LFQ) [4,5] yielding protein identifications from the normalised top 3 peptide intensities (Table S2). To confirm we could distinguish between OAC and NAT tissues using protein expression we performed Principal Component Analysis (PCA) using the normalised top 3 peptide intensities of the 500 most variable proteins. (Figure 1) [6]. The PCA yielded clear separation between OAC and NAT along PC1 accounting for 45% of the variance between the tissues. However, whilst the NAT samples were tightly grouped, the tumour samples were more dispersed, indicating some heterogeneity between tumours (Figure 1 A). Over 90% of the variation between the matched OAC and NAT samples is accounted for by the first 10 principal components (Figure 1 B).

**Figure 1:**
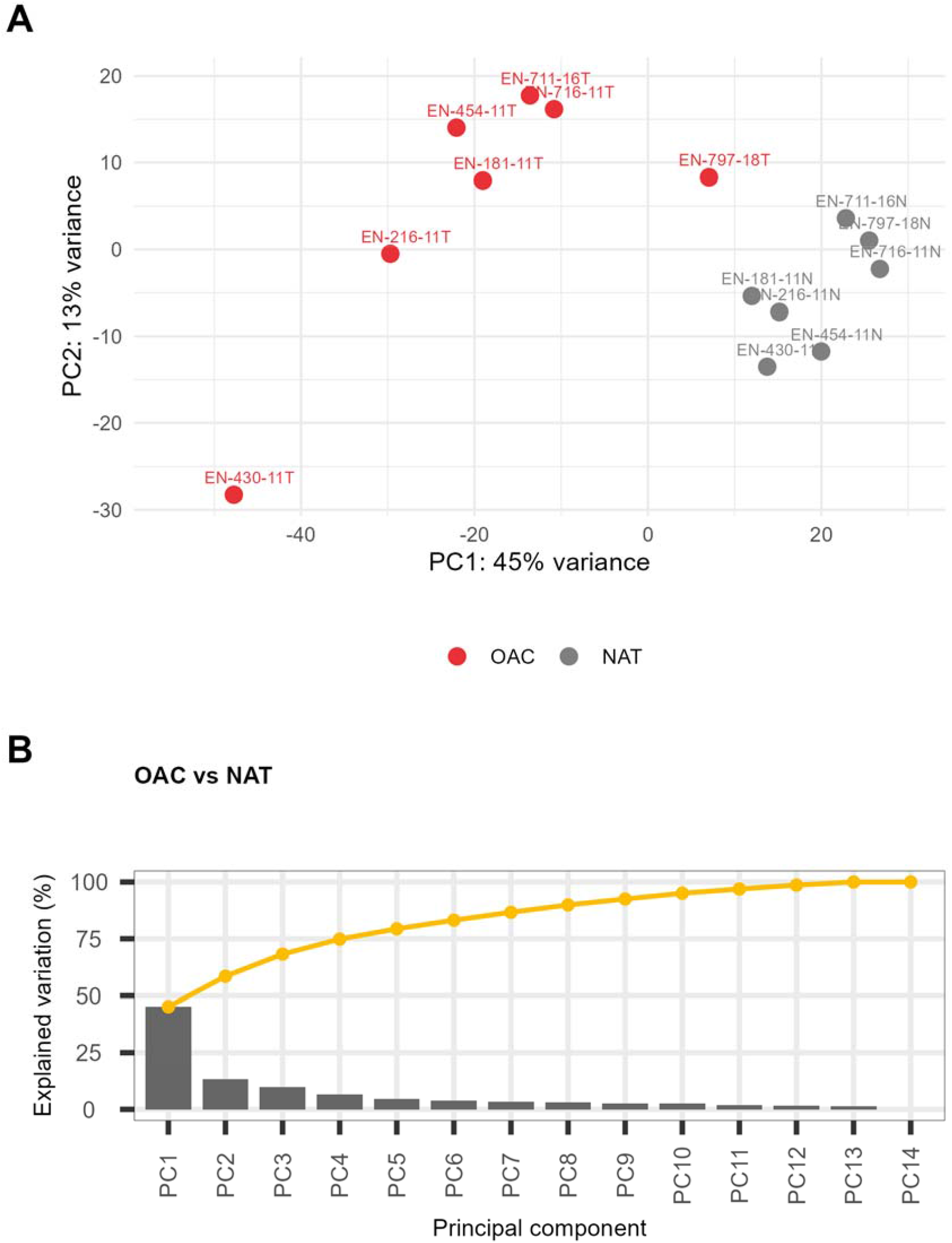
**Principal Component Analysis of OAC and NAT (A) PCA of normalised top 3 peptide intensities of 500 most variable proteins. OAC (red) & NAT (grey). Samples are numbered with donor identifier. (B) Scree plot of contribution of principal components to total variance.**

We then grouped the samples according to OAC or NAT and calculated differential protein expression (DEP) [7]. Of the 3552 proteins, we found 419 DEPs for OAC and 40 DEPs for NAT at thresholds of a log_2_ fold-change of greater than 1 and p-value of less than 1% (Figure 2 A). These and the other thresholds used here are necessarily arbitrary and chosen to balance being conservative whilst not over-excluding information. The data without thresholds is provided in Supporting Information Table S3. Anterior Gradient 2 (AGR2), involved protein folding and secretion was the most DEP for OAC. High AGR2 gene expression is a known unfavourable prognostic marker in renal and liver cancer [8]. AGR2 has also been identified as highly expressed protein in other cancers including OAC [9,10]. Amongst others and also as previously reported as putative immunohistological markers, we identified high expression in OAC of Endoplasmic reticulum chaperone BiP (HSPA5), Deoxynucleoside triphosphate triphosphohydrolase SAMHD1 (SAMHD1), Rho GDP-dissociation inhibitor 2 (ARHGDIB) ; and high expression in NAT of Protein-glutamine gamma-glutamyltransferase E (TGM3) and Heat shock protein beta-1 (HSPB1) [10,11].

**Figure 2:**
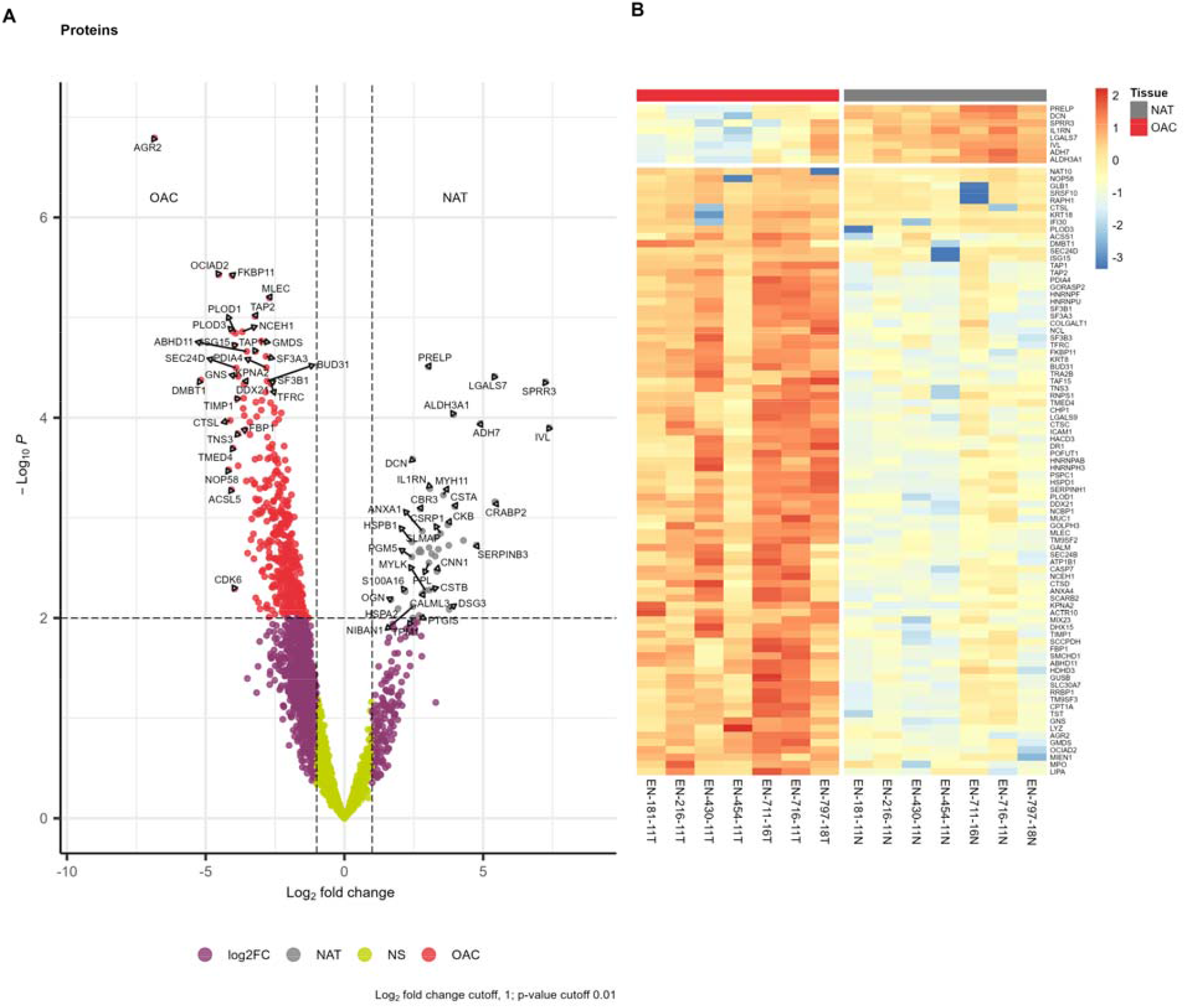
**(A) Volcano plot of differentially expressed proteins for OAC vs NAT. Proteins are labelled with gene names. Thresholds are represented by dotted lines at p-value of 1% and log_2_ fold change of 1. (B) Heatmap of DEPs below a FDR of 2% (n=92). Colour bar shows log_2_ fold change rescaled as z-scores i.e. each unit from zero represents one standard deviation from the row average value for each protein (C) Bar plots of enrichment of OAC GO Biological Processes. The top 25 pathways are shown with statistical significance level indicated by the -log_10_ p-value on the x-axis. Proteins were selected using thresholds for OAC DEPs above a log_2_ fold change 2 and below a FDR of 5% (n=232).**

Figure 2 B focuses in on the most statistically significant DEPs (92 proteins below FDR of 2%) across the OAC cohort. We found high expression for proteins in OAC relating to cell structure such as Keratin, type II cytoskeletal 8 (KRT8), RNA processing and protein folding such as Nucleolar RNA helicase 2 and (DDX21), Peptidyl-prolyl cis-trans isomerase (FKB11), and the immune system such as Antigen peptide transporters 1 & 2 (TAP1, TAP2) and Intercellular adhesion molecule 1 (ICAM1), suggesting that tumourigenesis may impact cellular phenotype and immunoregulation. A striking observation is for donor EN-454-11 and Nucleolar protein 58 (NOP58) which is generally highly expressed in OAC except for EN-454-11. We previously reported direct observation a putative neoantigen eluted from tumour HLA for EN-454-11 derived from mutation G95R in NOP58 [3]. Calculation of the Gibbs free energy of unfolding (ΔΔG) between the wild type and mutant NOP58 protein using DDGun [12] predicts the G95R mutation decreases the stability of NOP58 (Supporting Information).

Finally we performed functional analysis using 232 OAC DEPs (log_2_ fold-change greater than 2 and below FDR 5%). We identified greatest enrichment for pathways of biological processes relating to RNA processing, particularly mRNA splicing (Figure 3 A). For enrichment of Reactome pathways [13] also found changes in RNA processing, but also in immunological pathways, specifically Neutrophil degranulation and antigen processing and presentation (Figure 3 B).

**Figure 3:**
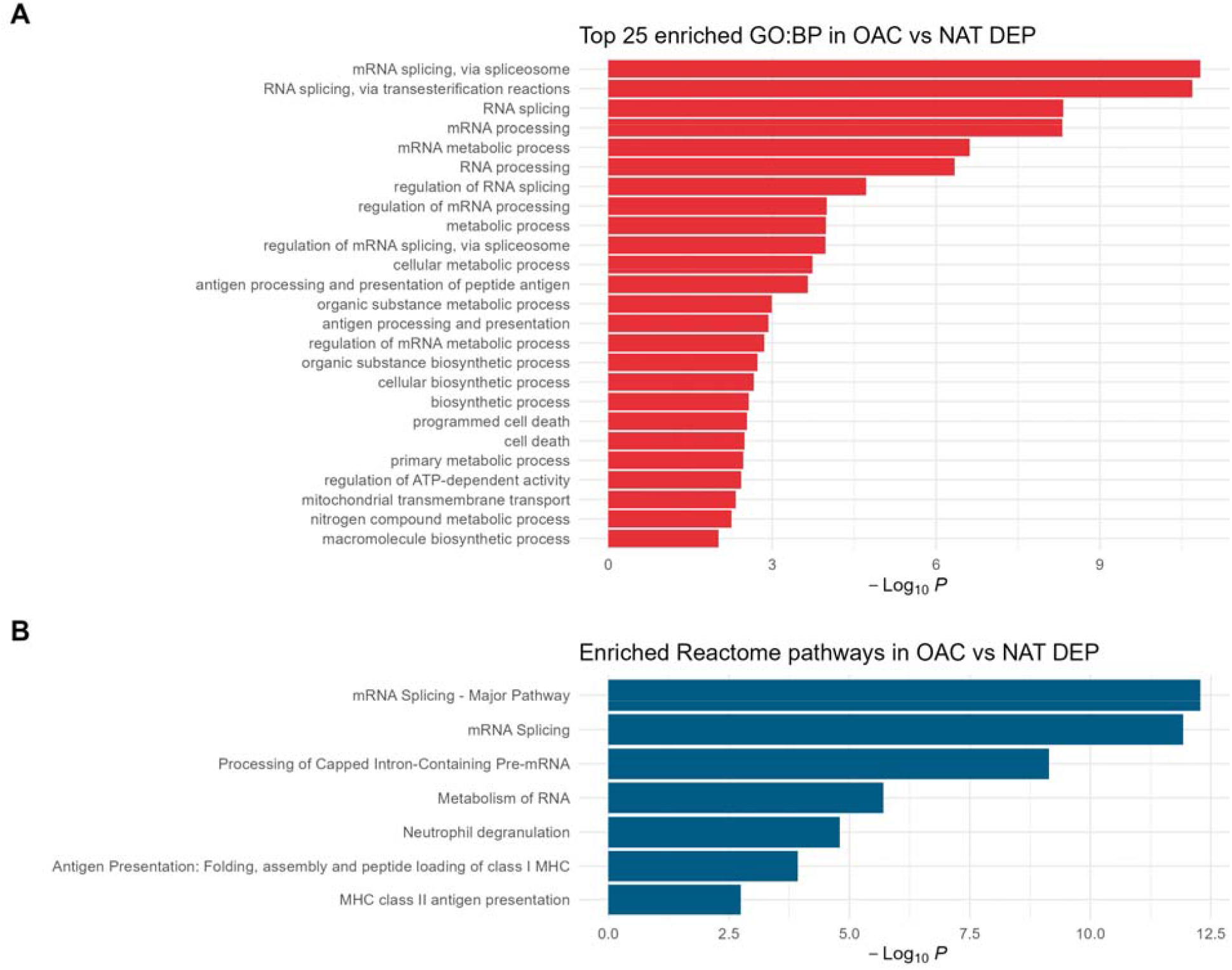
**Functional analysis bar plots of OAC (A) Enriched GO Biological Processes. The top 25 pathways are shown. (B) Enriched Reactome pathways. Statistical significance level indicated by the -log_10_ p-value on the x-axis. Proteins were selected using thresholds for OAC DEPs above a log_2_ fold change 2 and below a FDR of 5% (n=232).**

## Discussion

Previously we used proteogenomics analysis to identify patient specific neoantigens arising OAC mutations as therapeutic targets [3]. Here we compared the proteomes of OAC tissues to NAT from seven patients of the same cohort. This was a small study, limiting the extent to which our findings generalise. For example, epithelial cell adhesion molecule (EPCAM) was not uniformly expressed across all samples, so we are unable to confirm reports of high expression of EPCAM as a putative OAC biomarker [10,14]. The other main limitation in our design is that we are unable to compare these differential protein expression observations to other information that might support or discount their value, such as gene expression.

The principal value of the results presented here is the quantification of 3,500 proteins of which nearly 500 were differently expressed as a resource to other OAC researchers.

Additionally, the finding that the G95R mutation decreases the stability of NOP58 suggests that for donor EN-454-11, NOP58 is more likely to be a defective ribosomal product (DRiP) [15,16], which on the one hand leads to the decreased expression seen in Figure 2 B, whilst on the other increasing the probability of NOP58 neoantigen presentation. This raises the interesting potential of quantifying protein stability changes affected by single nucleotide polymorphisms and their effects on DRiP-derived neoantigen processing.

Our functional analysis corresponds with previous reports characterising changes in OAC tissues, and identifying targets for checkpoint inhibition such as G2/M checkpoint related protein Cell division cycle 5-like protein (CDC5L) [17]. Enrichment of the neutrophil degranulation pathway is indicative of inflammation, a known risk factor in cancer [18].

Overall, these observations offer independent corroboration and contrast to existing studies seeking to identify biomarkers or targets for more effective OAC specific treatments. Moreover our NOP58 observation suggests another parameter, protein stability, that can be used in the prediction of putative neoantigens for personalised therapies.

## Materials and Methods

### Tissue preparation

Subjects diagnosed with OAC were recruited to the study (see Table S1 for clinical characteristics). Tumours were excised from resected oesophageal tissue post-operatively by pathologists and processed either for histological evaluation of tumour type and stage, or snap frozen at −80°C.

### Protein extraction and digestion

Snap frozen tissue samples were briefly thawed and weighed prior to 30s of mechanical homogenization (Fisher, using disposable probes) in 4mL lysis buffer (0.02M Tris, 0.5% [w/v] IGEPAL, 0.25% [w/v] sodium deoxycholate, 0.15mM NaCl, 1mM ethylenediaminetetraacetic acid (EDTA), 0.2mM iodoacetamide supplemented with EDTA-free protease inhibitor mix). Homogenates were clarified for 10min at 2000g, 4°C and then for a further 60min at 13 500g, 4°C.

Protein concentration of tissue lysates was determined by BCA assay, and volumes equivalent to 100 mg of protein were precipitated using methanol/chloroform as previously described [19]. Pellets were briefly air-dried prior to resuspension in 6 M urea/50 mM Tris-HCl (pH 8.0). Proteins were reduced by the addition of 5 mM (final concentration) DTT and incubated at 37°C for 30 min, then alkylated by the addition of 15 mM (final concentration) iodoacetamide and incubated in the dark at room temperature for 30 min. 4 µg Trypsin/LysC mix (Promega) were added and the sample incubated for 4 h at 37°C, then 6 volumes of 50 mM Tris-HCl pH 8.0 were added to dilute the urea to < 1 M, and the sample was incubated for a further 16 h at 37°C. Digestion was terminated by the addition of 4 µL of TFA, and the sample clarified at 13,000 x g for 10 min at RT. The supernatant was collected and applied to Oasis Prime microelution HLB 96-well plates (Waters, UK) which had been pre-equilibrated with acetonitrile. Peptides were eluted with 50 µL of 70% acetonitrile and dried by vacuum centrifugation prior to resuspension in 0.1% formic acid.

### Mass spectrometry proteomics

8 µg of peptides per sample were separated by an Ultimate 3000 RSLC nano system (Thermo Scientific) using a PepMap C18 EASY-Spray LC column, 2 µm particle size, 75 µm x 75 cm column (Thermo Scientific) in buffer A (H_2_O/0.1% Formic acid) and coupled on-line to an Orbitrap Fusion Tribrid Mass Spectrometer (Thermo Fisher Scientific,UK) with a nano-electrospray ion source.

Peptides were eluted with a linear gradient of 3-30% buffer B (acetonitrile/0.1% formic acid) at a flow rate of 300 µL/min over 200 min. Full scans were acquired in the Orbitrap analyser in the scan range 300-1,500 m/z using the top speed data dependent mode, performing an MS scan every 3 second cycle, followed by higher energy collision-induced dissociation (HCD) MS/MS scans. MS spectra were acquired at a resolution of 120,000, RF lens 60% and an automatic gain control (AGC) ion target value of 4.0e5 for a maximum of 100 ms. MS/MS scans were performed in the ion trap, higher energy collisional dissociation (HCD) fragmentation was induced at an energy setting of 32% and an AGC ion target value of 5.0e3.

### Proteomic data analysis

Raw spectrum files were analysed using Peaks Studio 10.0 build 20190129 [4,20] and the data processed to generate reduced charge state and deisotoped precursor and associated product ion peak lists which were searched against the UniProt database (20,350 entries, 2020-04-07) plus the corresponding mutanome for each sample (∼1,000-5,000 sequences) and contaminants list in unspecific digest mode. Parent mass error tolerance was set a 10ppm and fragment mass error tolerance at 0.6 Da. Variable modifications were set for N-term acetylation (42.01 Da), methionine oxidation (15.99 Da), carboxyamidomethylation (57.02 Da) of cysteine. A maximum of three variable modifications per peptide was set. The false discovery rate (FDR) was estimated with decoy-fusion database searches [4] and were filtered to 1% FDR.

### Differential protein expression

Label free quantification using the Peaks Q module of Peaks Studio [4,5] yielding matrices of protein identifications as quantified by their normalised top 3 peptide intensities. The resulting matrices were filtered to remove any proteins for which there were more than two missing values across the samples. Differential protein expression was then calculated with DEqMS using the default parameters [7].

Principal component analysis of the normalised top 3 peptide intensities was performed using DESEq2 [6] and PCATools [21].

Results were visualised using EnhancedVolcano [22], pheatmap [23] and ggplot2 [24].

### Functional analysis

Functional enrichment analysis was performed using g:Profiler [25] using default settings for homo sapiens modified to exclude GO electronic annotations. Protein ids were used as inputs.

## Supporting Information

Supporting Information comprises of Tables S1-S4 as csv files, and the prediction of the effect of the G95R mutation on NOP58 using DDGun.

It is available on Github: https://github.com/ab604/oac-globals-supplement and Zenodo DOI: 10.5281/zenodo.13799179

### Data availability

The mass spectrometry proteomics data have been deposited to the ProteomeXchange Consortium via the PRIDE [26] partner repository with the dataset identifier PXD054428 and 10.6019/PXD054428.

Reviewer account details: Username: Password: o6YBxzmg0sn9

### Funding Information

This study was supported by a Cancer Research UK Centres Network Accelerator Award Grant (C328/A21998). Instrumentation in the Centre for Proteomic Research is supported by the BBSRC (BM/M012387/1).

## Ethics statement

Informed written consent was provided for participation by all individuals. Ethical approval for this study was granted by the Proportionate Review Sub-Committee of the North East – Newcastle & North Tyneside 1 Research Ethics Committee (Reference 18/NE/0234). This study was approved by the University of Southampton Research Ethics Committee.

